# Conserved domains and structural motifs that differentiate closely related Rex1 and Rex3 DEDDh Exoribonuclease families are required for their function in yeast

**DOI:** 10.1101/2024.04.22.590624

**Authors:** Peter W. Daniels, Sophie Kelly, Iwan J. Tebbs, Phil Mitchell

## Abstract

The DEDD family of exonucleases has expanded through evolution whilst retaining a well conserved catalytic domain. One subgroup of DEDD exoribonucleases with very closely related catalytic domain sequences includes the yeast enzymes Rex1 (RNA exonuclease 1) and Rex3, the metazoan REXO1 (RNA exonuclease 1 homologue) and Rexo5 proteins, and the plant protein Sdn5 (small RNA degrading nuclease). Comparison of Alphafold models for these proteins reveals that Rex1, Rexo5 and Sdn5 are structural homologues, consisting of a central catalytic domain inserted within a discontinuous alkaline phosphatase (AlkP) domain. The core AlkP domain of Rex1-related proteins contains distinct surface insertions in different eukaryotic lineages. Yeast Rex1 contains three loops that are modelled to be directed towards the DEDD domain, one of which forms an extended helical arch that is conserved across fungi and plants but absent in metazoan homologues. We show that the arch and an adjacent loop are required for Rex1-mediated processing of 5S rRNA and tRNA in *Saccharomyces cerevisiae*. Rex3 structural homologues, including REXO1, contain a KIX domain (CREB kinase-inducible domain (KID) interacting domain) and a cysteine- and histidine-rich domain (CHORD) adjacent to a C-terminal DEDD domain. In contrast to Rex1, Rex3 proteins are found in metazoans and fungi but not in plants or algae. Deletion of the KIX domain within yeast Rex3 blocked its function in RNase MRP processing. Taken together, this work identifies evolutionarily conserved structural hallmarks of Rex1 and Rex3 enzymes and demonstrates that these features are required for Rex1- and Rex3-mediated RNA processing pathways *in vivo*.

## Introduction

The DEDD family of 3’ exonucleases is a group of diverse enzymes that hydrolytically remove (deoxy)ribonucleotide 5’-monophosphates from the 3’ end of DNA or RNA. They are found throughout biology and have a broad range of actions in the key biological processes of DNA replication and repair, gene expression and RNA turnover. The catalytic centres of these enzymes contain four conserved acidic residues (hence the term DEDD) that are located within three conserved sequence motifs known as the Exo I, Exo II and Exo III motifs (Zuo and Deutscher 2001). The DEDD family is split into DEDDh and DEDDy subfamilies that contain either a conserved histidine or tyrosine residue within the Exo III motif. The DEDD catalytic domain is found in both exonucleases and in polymerases that have associated proof-reading activities (Moser et al. 1997; Zuo and Deutscher 2001).

While the *Escherichia coli* genome encodes two DEDDh family exoribonucleases, there are eight encoded within the genome of the budding yeast *Saccharomyces cerevisiae* and fourteen expressed in human cells. These enzymes show specificity for some substrates while functioning redundantly in the processing or degradation of others. For example, the yeast enzymes Rex1-3 have overlapping yet distinct roles in stable RNA processing (van Hoof et al. 2000). The RNase T functional homologue Rex1 (RNA exonuclease 1) has specific roles in the 3’ end maturation of 5S rRNA, 25S rRNA, signal recognition particle (SRP) RNA and some tRNAs (Copela et al. 2008; Foretek et al. 2016; Kempers-Veenstra et al. 1986; Leung et al. 2014; Ozanick et al. 2009; Piper et al. 1983; van Hoof et al. 2000). Rex3 is specifically required for correct 3’ end processing of the RNA subunit of RNase MRP (van Hoof et al. 2000) and functions together with Rex2 in the autoregulated degradation of *RTR1* mRNA transcripts (Hodko et al. 2016). None of the yeast DEDDh exonucleases are essential for mitotic growth but *rex1* null alleles are synthetic lethal with null alleles of the *RRP6* gene that encodes a DEDDy family exoribonuclease (Garland et al. 2013; van Hoof et al. 2000) or the *RRP47* gene (Peng et al. 2003) that encodes an Rrp6-associated protein (Mitchell et al. 2003).

rRNA processing events that are mediated by RNase T in *E. coli* and Rex1 in *S. cerevisiae* are carried out by the DEDDh family enzyme Rexo5 in *Drosophila melanogaster* (Gerstberger et al. 2017). Humans and mice express a Rexo5 homologous protein (known as NEF-sp or REXO5) that was characterised as a nuclease whose expression is highly enriched in the testes (Silva et al. 2017). Whether the human NEF-sp/REXO5 protein functions in 5S and 28S rRNA 3’ processing in either germline or somatic cells is currently unclear. Human cells express another Rex1-related, DEDDh family enzyme known as RNA exonuclease 1 homologue (REXO1). REXO1 was identified as a protein that interacts directly with the transcription elongation factors Elongin A and SII (TFIIS) and was recognised to have a C-terminal domain with homology to RNase T (Tamura et al. 2003). Beyond this sequence homology, the relationship between yeast Rex1 and REXO1 has not been addressed. The SDN (Small RNA degrading nuclease) gene family in *Arabidopsis thaliana* was identified through sequence homology with the yeast Rex proteins (Ramachandran and Chen 2008). Genetic depletion of Sdn1, Sdn2 and Sdn3 led to defects in miRNA processing. We have previously reported that Sdn5 is structurally related to Rex1 (Daniels et al. 2022).

Yeast Rex1 has a molecular weight of 63 kDa, approximately twice that of Rex2 or Rex4. A high confidence Alphafold model suggests that Rex1 contains a discrete alkaline phosphatase (AlkP) domain, in addition to the catalytic DEDDh domain (Daniels et al. 2022). The AlkP domain of Rex1 is required for its stable expression *in vivo* and for its catalytic activity *in vitro* (Daniels et al. 2022) but structural features within the AlkP domain of Rex1 that are required for its function have not yet been determined.

We have carried out phylogenetic sequence analyses of the very closely related Rex1 and Rex3 exonucleases and compared predicted structural models for these proteins, with the aim of identifying structural characteristics of each enzyme. We report that the AlkP domain is a common feature of homologous Rex1, Rexo5 and Sdn5 proteins that are found in all major eukaryotic lineages. Mutagenesis studies reveal that insertions on the surface of the AlkP domain of Rex1, including the highly conserved helical “arch” that is modelled to be directed towards the DEDDh domain, are required for 5S rRNA and tRNA processing in *S. cerevisiae*. Furthermore, we show that REXO1 and Rex3 proteins have a common domain architecture, consisting of a conserved KIX (CREB kinase-inducible domain (KID) interacting domain) domain and a single CHORD (cysteine- and histidine-rich domain) domain (Heise et al. 2007; Shirasu et al. 1999; Thakur et al. 2014), in addition to the DEDDh domain. Deletion analyses show that the N-terminal KIX domain within yeast Rex3 is required for its function in the 3’ end maturation of RNase MRP RNA *in vivo*. This study identifies structural hallmarks of Rex1 and Rex3 exoribonucleases that are required for their function in specific RNA processing pathways.

## Results and Discussion

### The AlkP domain is conserved in Rex1-related proteins

In contrast to the other Rex proteins in yeast, the catalytic domain of Rex1 lies in the central region of the protein, positioned between comparably long N- and C-terminal regions. Alphafold (Jumper et al. 2021; Varadi et al. 2022) predicts a high confidence dual domain structure for yeast Rex1 (a mean pLDDT score of 92 for residues 53-553) consisting of a central DEDDh catalytic domain that is inserted within a domain comprising sequences from both the N- and C-terminal regions of the protein (Daniels et al. 2022). This discontinuous domain has the same fold as experimentally determined structures for phosphopentomutases (PPMs) and cofactor-independent phosphoglycerate mutases (iPGMs) (Jedrzejas et al. 2000; Panosian et al. 2011), two members of the alkaline phosphatase (AlkP) superfamily (Galperin and Jedrzejas 2001) (Fig. 1). Here, we refer to this domain as the AlkP domain of Rex1.

**Figure 1.**
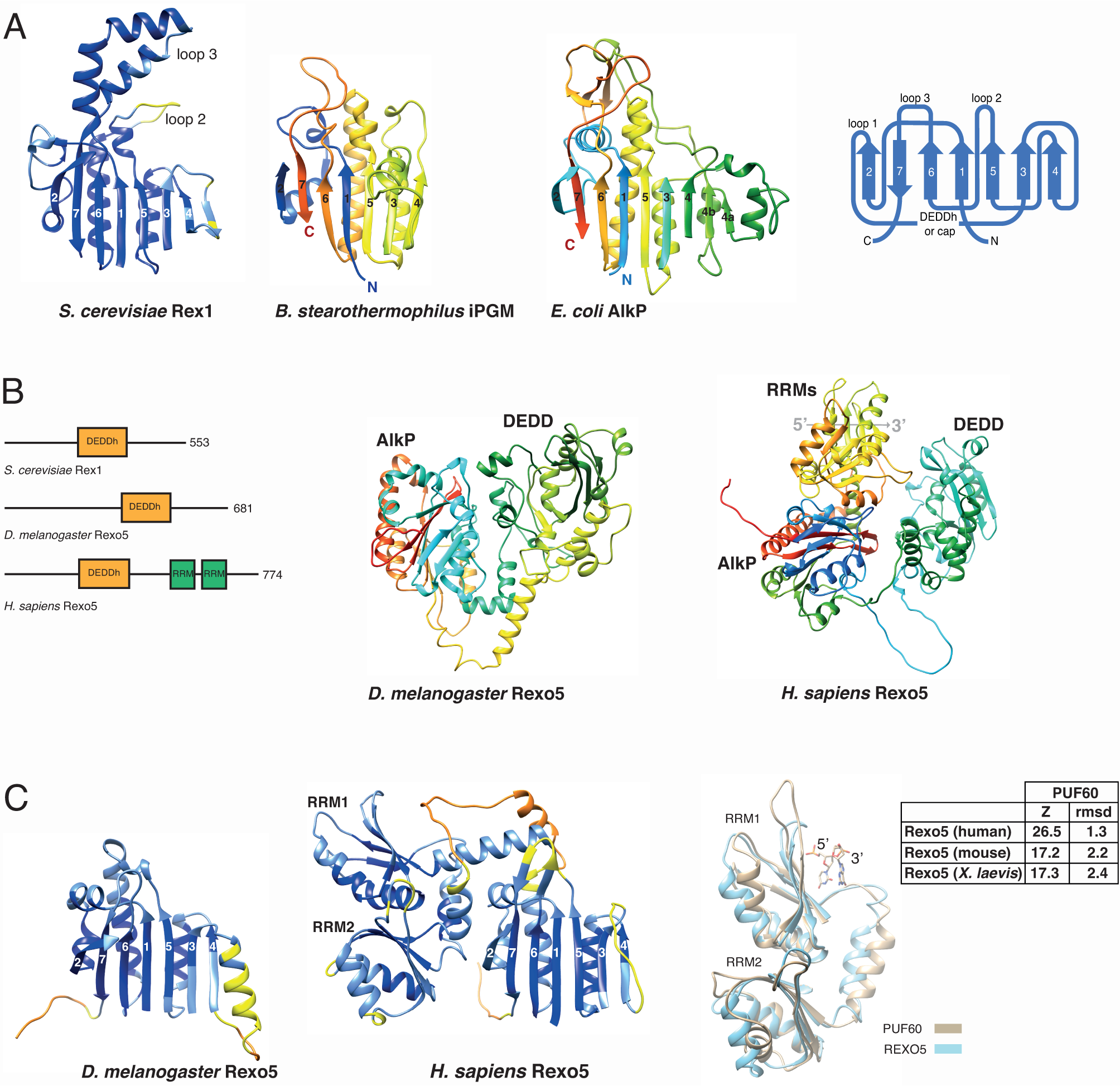
Rex1 and Rexo5 orthologues have a discontinuous alkaline phosphatase fold. (A) Ribbon structures of AlkP and the AlkP domains in Rex1 and iPGM, together with a schematic of the core domain. The β strands are numbered and prominent loops in Rex1 are labelled. Regions between strands 2 and 3 (and connecting strands 3 and 4 within AlkP) are omitted for clarity. The Rex1 model is coloured to reflect pLDDT confidence scores (>90%, dark blue; >70%, light blue; >50%, yellow; <50%, orange). iPGM and AlkP models are rainbow-coloured (blue to red, N’ to C’). (B) Domain structure of Rex1 and Rexo5 proteins, and rainbow-coloured ribbon structure models of the fly and human Rexo5 proteins (residues 145-681 and 49-774, respectively). The 5’-3’ orientation of bound RNA across the surface of RRM1 in the model of the human protein is indicated. (C) Models of the Rexo5 AlkP domains, coloured according to pLDDT scores and viewed from the same perspective as yeast Rex1 in (A), and a structural overlay of the twin RRM domains in human Rexo5 and PUF60 with bound dinucleotide. Z-scores and rmsd values for PUF60 structure overlays with vertebrate REXO5 proteins are shown.

The core AlkP domain fold consists of a predominantly parallel, seven stranded β-sheet that is supported by α-helices on both sides (Fig. 1A, helices and loops on the internal face are not shown for clarity). The β-strands are numbered 1-7 in figure 1, from the N- to C-terminus. The central DEDDh domain of Rex1 is connected to the AlkP domain by flexible polypeptide linkers that lead from β2 into the DEDDh domain, and from the DEDDh domain back into β3. Prominent surface loops of the yeast Rex1 structure are referred to here as loop 1, that leads from β2 into the linker connecting the AlkP and DEDDh domains, loop 2 that connects β5 and β6, and loop 3 that connects β6 and β7 (Fig. 1A).

Analysis of structure similarity clusters within Alphafold (Barrio-Hernandez et al. 2023; Steinegger and Soding 2018) revealed that the dual domain architecture of yeast Rex1 is found in proteins from fungi, flowering plants (including several Sdn5 homologues), green algae, amoebozoa, diatoms and flagellates, as well as in the metazoan organisms *Clytia hemisphaerica* (a cnidarian) and *Hirondella gigas* (an amphipod). The structure similarity clusters do not include the functionally homologous Rexo5 protein from *D. melanogaster* or the vertebrate homologues NEF- sp/REXO5 or REXO1, although reverse analysis of structure similarity clusters of the *D. melanogaster* Rexo5 protein identified Rex1 homologues in other yeasts. The DEDDh domain of Rexo5, like Rex1, is also positioned between extensive N- and C-terminal flanking regions. Vertebrate REXO5 proteins also have a central DEDDh domain and contain two RNA recognition motif (RRM) domains within their C-terminal regions that are not seen in Rexo5 from *D. melanogaster* or yeast Rex1 (Gerstberger et al. 2017; Silva et al. 2017) (Fig. 1B). Comparison of high confidence Alphafold models for Rexo5 proteins from *D. melanogaster* and humans (mean pLDDT scores of 84 and 80 for the residues shown in Fig. 1B, respectively) revealed the same DEDDh/AlkP dual domain structure seen in yeast Rex1. Structure comparisons using the Dali server revealed that the modelled AlkP domains of the fly and human proteins show high Z-scores and rmsd values with experimentally determined structures of AlkP domains (12.6 and 2.6Å, and 13.0 and 2.2Å, respectively, with the structure of iPGM from *Bacillus stearothermophilus* (Jedrzejas et al. 2000)). There is a low level of packing between the DEDDh and AlkP domains in the fly, human and yeast proteins, leading to variation in the mutual orientation of these domains in the models.

### Insertions within the AlkP domain of Rex1-related proteins

Non-conserved sequences within the alkaline phosphatase superfamily map as insertions on the surface of the common α/β fold (Galperin and Jedrzejas 2001). The central “cap” domain of PPMs and iPGMs is inserted between β2 and β3 of the AlkP domain (Jedrzejas et al. 2000; Panosian et al. 2011). Strikingly, the DEDDh domain within the dual domain structure of Rex1-related proteins is also inserted between β2 and β3. This common insertion point is consistent with observations that the interface between the AlkP domain and the cap or DEDDh domain may have functional significance. Both the AlkP and cap domains contribute to the catalytic centres of PPMs and iPGMs (Jedrzejas et al. 2000; Panosian et al. 2011). Furthermore, *in vivo* crosslinking experiments and structural threading studies suggest that both the DEDDh and AlkP domain of yeast Rex1 contribute to substrate binding (Daniels et al. 2022).

The relative size and complexity of the loop insertions varies across Rex1-related proteins. Loop 3 is extended in yeast and plant proteins but contracted to a short loop in Rexo5 from *D. melanogaster* and humans. Loop 2 is also contracted in the fly protein but expanded within the vertebrate proteins to include the RRMs (Fig. 1C). We observed that Rex1-related proteins from trypanosomes are predicted to have an additional extended loop between strands 1 and 2.

The predicted structures of the two RRMs within vertebrate Rexo5 proteins have very high similarities to the twin RRMs that constitute the *half pint* domain (Dali score of 26.5-17.2, rmsd of 2.4-1.3 Å) found within PUF60/FIR proteins (Van Buskirk and Schupbach 2002) (Fig. 1C). Furthermore, models of the human and mouse Rexo5 proteins were the most similar structures identified in searches comparing the PUF60 RRM structure (5KWQ) against species-specific Alphafold databases. The N-terminal RRM of PUF60/FIR (labelled RRM1 in Fig. 1C) is packed against the C-terminal RRM, obstructing its RNA binding surface (Crichlow et al. 2008; Hsiao et al. 2020). This packed alignment is dependent upon a long α-helix connecting the two RRMs (Crichlow et al. 2008) that is also predicted to be present within Rexo5 structures (Fig. 1C). The low predicted aligned error within the Alphafold model of human Rexo5 for the C-terminal region of the protein is consistent with a tight packing between the twin RRM domains and a close alignment of the RRMs with the AlkP domain. The orientation of bound nucleic acid on consensus RRMs is conserved (Daubner et al. 2013). Despite the low degree of packing between the DEDDh and AlkP domains, the Alphafold model of human Rexo5 is consistent with bound RNA passing across the surface of the N-terminal RRM towards the active site of the DEDDh domain in a 5’-3’ direction (Fig. 1B,C).

### Loops at the domain interface of Rex1 are required for 5S rRNA and tRNA processing

A striking feature of the yeast Rex1 model is that the three prominent loops on the surface of the AlkP domain are projected towards the catalytic DEDDh domain (Fig. 2A). Loop 3 forms an extended arch structure and is clearly identifiable in the Alphafold structural similarity clusters of Rex1-related proteins from fungi, amoebozoa, diatoms, plants and green algae. We replaced loops 1-3 within Rex1 from *S. cerevisiae* with short linker peptides and assayed the ability of the deletion alleles to complement the synthetic lethal growth phenotype of a *rex1Δ rrp47Δ* double mutant and the 5S rRNA and tRNA processing defects observed in a *rex1Δ* mutant. Wild-type and mutant Rex1 proteins were expressed as N-terminal fusions to assess effects of the mutations on protein expression levels.

**Figure 2.**
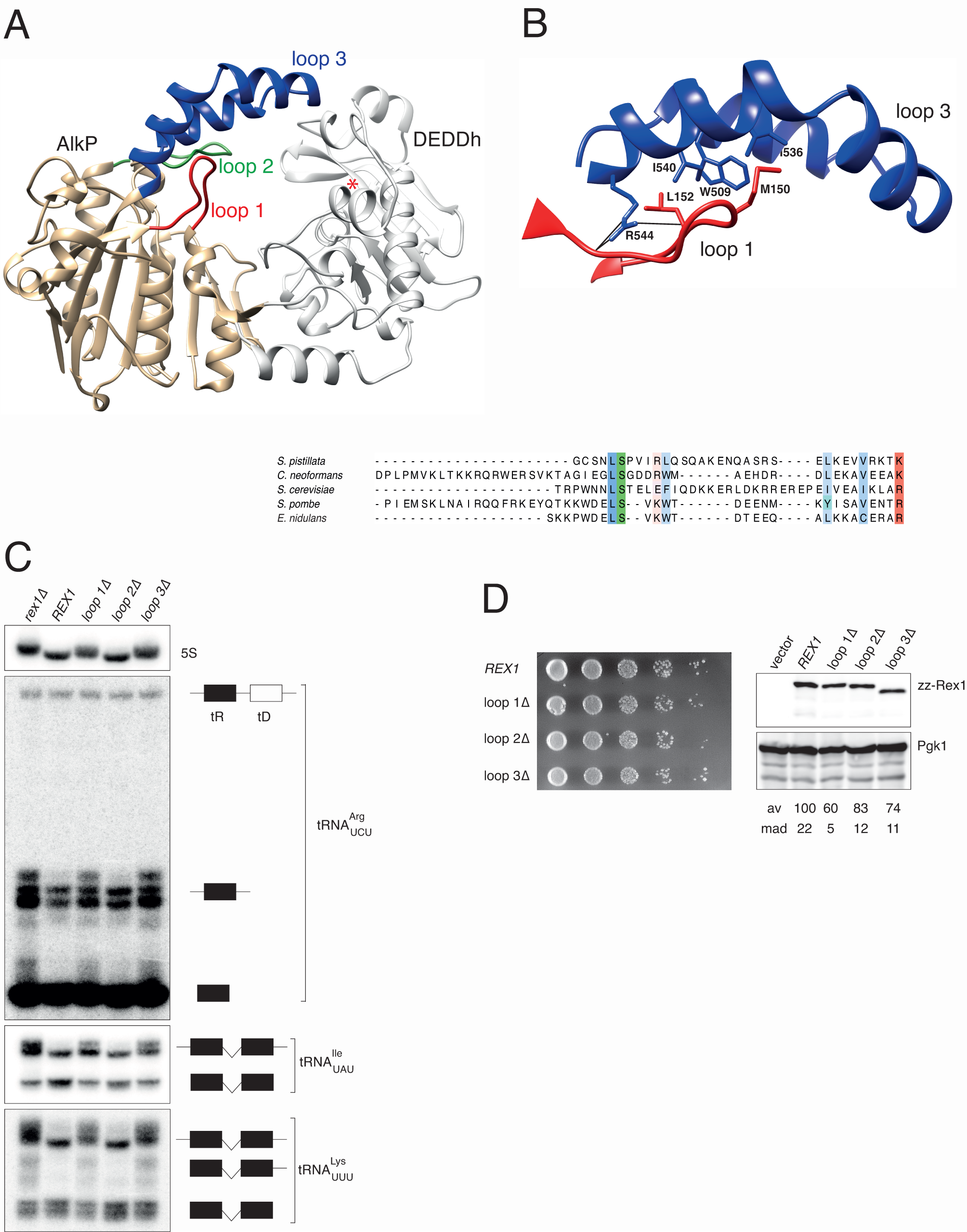
Loops within the AlkP domain of Rex1 are required for RNA Processing *in vivo*. (A) Alphafold model of yeast Rex1 (residues 53-553), showing the position of surface loops on the AlkP domain. (B) Expanded view of loops 1 and 3, and an alignment of loop 1 sequences from high confidence Alphafold models of Rex1 proteins. (C) Northern blot analyses of RNA from a *rex1Δ* mutant and isogenic transformants expressing wild-type or mutant Rex1 proteins. Membranes were probed with oligonucleotides complementary to 5S rRNA, tRNA^Arg^_UCU_ and intronic sequences within tRNA^Ile^_UAU_ or tRNA^Lys^_UUU_. Detected tRNA species inferred from published data (Copela et al. 2008; Foretek et al. 2016; van Hoof et al. 2000) are indicated schematically on the right. (D) Spot growth assay of *rex1 rrp47Δ* double mutants and relative expression levels of the *rex1* mutant proteins. Western blot analyses were performed for wild-type and mutant *rex1* proteins expressed in a *rex1Δ* single mutant. Rex1 expression levels were normalised to Pgk1. Values for the mean average (av) and mean absolute deviation (mad) of three independent replicates are given.

The wild-type *REX1* allele and each loop deletion mutant supported growth of the *rex1Δ rrp47Δ* double mutant in plasmid shuffle assays. Furthermore, FOA-resistant *rex1 rrp47Δ* isolates expressing the loop mutants showed comparable growth to isolates expressing the *REX1* wild-type allele (Fig. 2D). The expression levels of the mutants were ∼ 60-80% of the expression level of the wild-type protein. This reduction is not predicted to be limiting for Rex1 function, since expression of plasmid-borne fusion proteins is more than three times higher than the expression level of a fully functional, chromosomally encoded fusion protein (Daniels et al. 2022).

Yeast 5S rRNA is extended at the 3’ end by 3-4 nucleotides in the *rex1Δ* mutant, compared to a wild-type strain (Piper et al. 1983; van Hoof et al. 2000). Strains lacking Rex1 are also defective in the 3’ end maturation of some tRNAs, including tRNA^Arg^_UCU_, tRNA^Ile^_UAU_ and tRNA^Lys^_UUU_ (Copela et al. 2008; Foretek et al. 2016; Piper and Straby 1989; van Hoof et al. 2000). Some tRNA^Arg^_UCU_ genes are expressed as the 5’ cistron of dicistronic tRNA transcripts including tRNA^Asp^_GUC_. Processing of tRNA^Arg^_UCU_ from the dicistronic transcript involves release of a ∼ 92 nucleotide long, 5’- and 3’ extended intermediate that is subsequently processed to the mature tRNA (Engelke et al. 1985). Northern blot hybridisation using a probe complementary to both the 3’ end of tRNA^Arg^_UCU_ and the downstream trailer sequence detected, under our hybridisation conditions, species with mobilities corresponding to the dicistronic primary transcript, 5’- and 3’-extended intermediates and the mature tRNA (Fig. 2C). Consistent with earlier reports (Piper and Straby 1989; van Hoof et al. 2000), extended forms of the processing intermediate and the mature tRNA were seen in the *rex1Δ* mutant. Probes specific to intronic sequences within tRNA^Ile^ and tRNA^Lys^ detected two major transcripts in wild-type cells. By comparison with published data (Copela et al. 2008; Foretek et al. 2016), these correspond to the primary transcript and shorter, end-processed but unspliced intermediates. The relative abundance of the primary transcript increased in the *rex1Δ* mutant, compared to the *REX1* wild-type strain, and the mutant accumulated extended forms of the primary transcript. Furthermore, tRNA^Lys^_UUU_ intermediates were observed in the *rex1Δ* mutant that were longer than the unspliced, end-processed form but shorter than the primary transcript. These observations are consistent with data from earlier studies showing a role for Rex1 in the 3’ end processing of tRNA^Arg^_UCU_, tRNA^Ile^ and tRNA^Lys^_UUU_ (Copela et al. 2008; Foretek et al. 2016).

Northern blot analyses showed that the 5S rRNA and tRNA processing defects seen in the *rex1Δ* mutant were complemented upon expression of the wild-type *REX1* allele or the loop 2Δ mutant but not in strains expressing the loop 1Δ or loop 3Δ mutant (Fig. 2C). These data show that loop 1 and loop 3 are required for Rex1-mediated 5S rRNA and tRNA processing but not for growth in a Rex1-dependent strain. These observations are not inconsistent, as the defect(s) in RNA processing and/or turnover underlying the synthetic lethal growth phenotype of *rex1Δ rrp47Δ* strains is not related to 5S rRNA or tRNA processing (Copela et al. 2008; Garland et al. 2013).

Loop 1 and loop 3 residues within the Alphafold model of yeast Rex1 have very high confidence mean pLDDT scores (93 for both polypeptide sequences) and are packed closely together to form an integrated structural module (Fig. 2B). Structure-based sequence alignment of loop 3 sequences based on high confidence Alphafold models (minimum mean pLDDT score of 81 for the sequences shown in Fig. 2) suggests that the packing of loops 1 and 3 in Rex1 is broadly conserved across fungi. Given the location of the loop 1/3 structural module at the domain interface, we speculate that the loop 1/3 structural module of yeast Rex1 contributes to substrate binding.

### Rex3 and REXO1 proteins share a common domain architecture

The catalytic domain of Rex1 is closely related at the sequence level to that of Rex3 (Gerstberger et al. 2017). In contrast to Rex1, but similar to Rex2 and Rex4, the DEDDh domain of Rex3 is found at the C-terminus of the protein. Structural overlay of Alphafold models for yeast Rex1 and Rex3 (mean pLDDT score of 88 for the entire length of the protein) suggested two prominent features of Rex3 that are not found in Rex1-related proteins (Fig. 3A). Firstly, the N-terminal region of yeast Rex3 is predicted to form a compact triple helix. Structure comparisons revealed a high degree of similarity between this predicted domain and the KIX domain (CREB kinase-inducible domain (KID) interacting domain) of the mediator subunit Med15/Gal11 of *Candida glabrata* and mouse CREB-binding protein (CBP) (Fig. 3B). Secondly, a ∼ 70 residue long, extended domain comprising three pairs of short antiparallel β-sheets is predicted to be positioned immediately upstream of the catalytic DEDDh domain. Structural homology searches revealed a common fold between this region and the cysteine- and histidine-rich domain (CHORD) of the Hsp90 cochaperone Rar1 (Zhang et al. 2010) (Dali Z-score of 4, rmsd of 3.4 Å).

**Figure 3.**
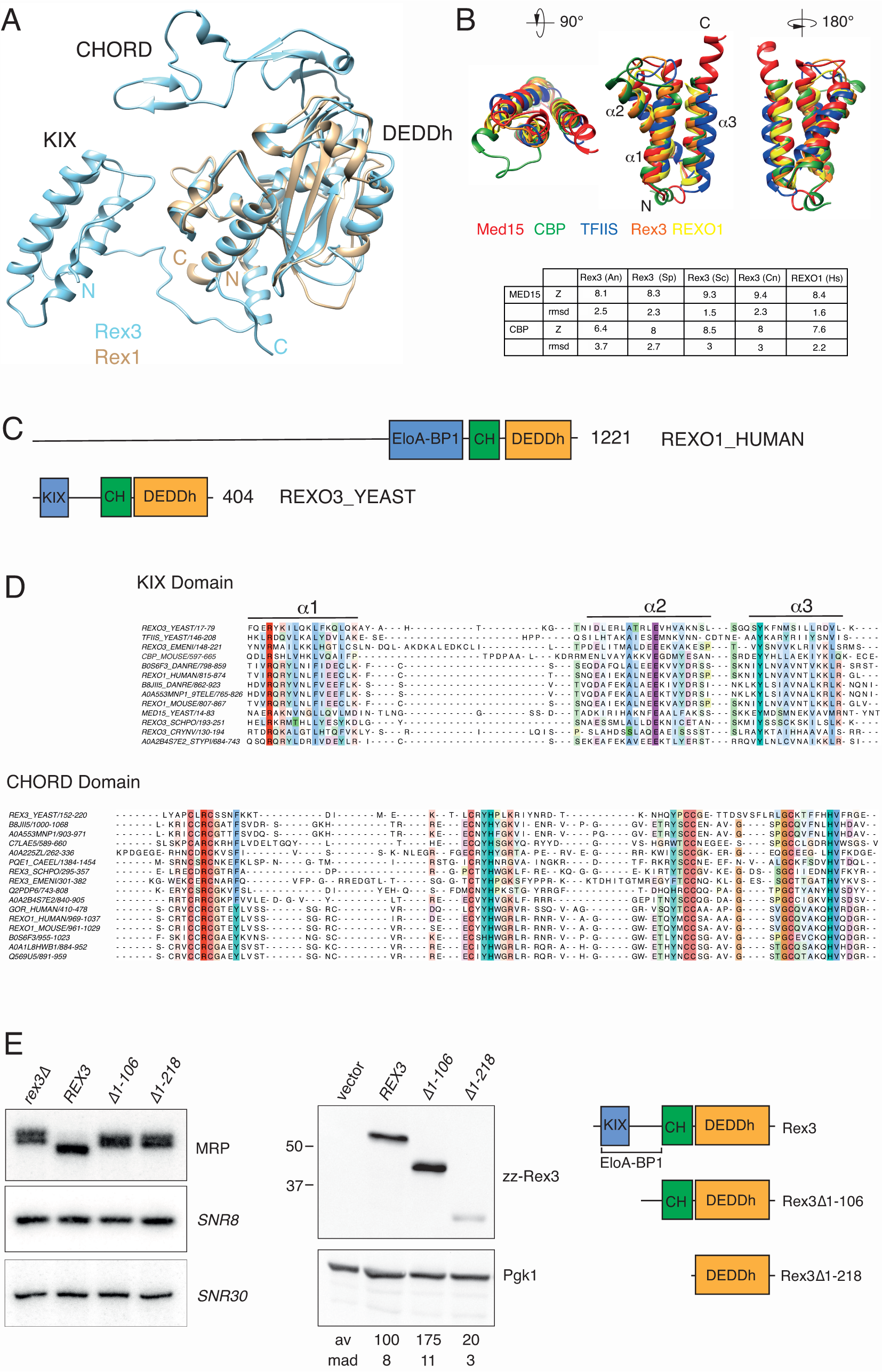
The KIX domain of Rex3 is required for MRP RNA Processing. (A) Structure overlay of Alphafold models for full-length Rex3 (blue) and the DEDDh domain of Rex1 (brown) from *S. cerevisiae*. Protein domains and termini are labelled. (B) Overlay ribbon diagrams of the triple helical domains within yeast Rex3 and human REXO1 proteins and the KIX domains of yeast Med15, mouse CBP and yeast TFIIS. Z-scores and rmsd values for the correlation between Rex3/REXO1 protein structural models and the structures of the KIX domains of Med15 and CBP are given. (C) Domain structure of human REXO1 and yeast Rex3 proteins. (D) Structure-based multiple sequence alignments of the KIX and CHORD domains of Rex3 and REOX1 proteins. Residues are coloured by conservation. The KIX domain alignment includes TFIIS, Med15 and CBP. (E) Analysis of RNA processing defects and protein expression levels of *rex3* mutants. Northern blot analyses were performed on RNA from a *rex3Δ* mutant and isogenic transformants expressing wild-type or mutant *rex3* alleles, using probes complementary to MRP RNA, snR8 and snR30. Western analyses were carried out on whole cell lysates from strains expressing zz-Rex3 fusion proteins. Rex3 expression levels, normalised to Pgk1 and expressed as a percentage of the wild-type protein, are indicated. Mean average (av) and mean absolute deviation (mad) values of three independent replicates are given. The schematic shows the domain architecture of the Rex3 proteins analysed.

These two structural features are also predicted within Alphafold models of Rex3 homologues from other yeasts, including *Aspergillus (Emericella) nidulans* and *Schizosaccharomyces pombe*. Moreover, the same domains are found at high confidence levels in the Alphafold model of the human protein REXO1, although the overall mean pLDDT score of the model is low. The central region of REXO1 that was shown to bind to Elongin A and TFIIS (Tamura et al. 2003) is annotated in Pfam as the Elongin A binding-protein 1 (EloA-BP1) domain (PF15870) and in InterPro as RNA exonuclease 1 homologue-like domain (REXO1-like domain (IPR031736). The EloA-BP1 domain is not only found in vertebrate and invertebrate REXO1 proteins but is also annotated for Rex3 proteins from the yeasts *S. pombe*, *Kluyveromyces lactis*, *Yarrowia lipolytica* and *Cryptococcus neoformans*. The annotated EloA-BP1 domain spans the helix bundle of the KIX domain and extends up to the CHORD domain, including a short helical stretch (residues 119-142 in yeast Rex3) that is packed against the DEDDh domain and conserved between Rex1 and Rex3 proteins.

The KIX domain within Med15 and CBP enables kinase-responsive protein interactions with disordered regions of transcription factors (Thakur et al. 2014). Other proteins that contain KIX domains include the DNA helicase RECQL5 and the transcription elongation factor TFIIS, both of which have been shown to interact directly with RNA polymerase II (RNAP II) (Kassube et al. 2013). Comparison of the helical domain with the Alphafold models for Rex3 and REXO1 proteins with the KIX domain of Med15, CBP and TFIIS revealed a high degree of structural similarity (Fig. 3B). The helices of the KIX domain are numbered 1-3 in Figure 3, from the N- to C-terminus. The structural fold of characterised KIX domains is dependent upon a conserved cation-π interaction between an arginine at the base of helix 1 and a tyrosine at the base of helix 3 (Thakur et al. 2014). These residues are conserved within the helical domains seen in Rex3- and REXO1-related proteins (Fig. 3D). Furthermore, a glutamate residue within helix 2 that is modelled to interact with both the arginine and tyrosine residues is also conserved across all KIX and EloA-BP1 domains.

Organism-specific Alphafold homology searches to identify proteins with domains of similar structure to that of the KIX domain of Med15 identified Rex3 proteins within *S. cerevisiae* and *S. pombe* as the highest hit, *D. melanogaster* REXO1 to be the second highest after RECQ5, and the human proteins REXO1 and GOR to have the highest homology after RECQ5, CBP and its paralogue P300. We conclude that the helical region within the EloA-BP1 domain seen in REXO1 and Rex3 proteins is structurally homologous to the KIX domain.

Interaction between RECQL5 and RNAP II leads to repression of transcription elongation. This is due, in part, to a competition between the KIX domains within RECQL5 and TFIIS for binding to the jaw domain of the Rpb1 subunit of RNAP II (Kassube et al. 2013). One possibility is that the KIX domain within REXO1 and Rex3 proteins may also compete with TFIIS for interaction with RNAP II and promote coupling of transcription termination with subsequent 3’ processing or degradation of the transcript. Notably, RNAP II is enriched with Rex3 in yeast mutants that are depleted of the transcription elongation factor Spt6 (Gopalakrishnan and Winston 2021).

CHORD domains comprise of two C_3_H motifs, containing three cysteines and a histidine residue, that each bind a Zn^2+^ ion (Heise et al. 2007; Shirasu et al. 1999). They mediate specific protein-protein interactions, notably with Hsp90 and the cochaperone protein Sgt1 (Shirasu et al. 1999; Takahashi et al. 2003; Zhang et al. 2010). The cysteine and histidine residues are conserved throughout the CHORD domain in Rex3-related proteins, as is their relative spacing (Fig. 3D), strongly suggesting that Rex3 and REXO1 are zinc-binding proteins. The CHORD domains of Rex3-related proteins are unusual inasmuch as they are single domains; CHORD domains are typically observed as tandem domains. CHORD domains are found in proteins from metazoans, plants and *Toxoplasma* but, notably, were previously reported as being absent from *S. cerevisiae* (Shirasu et al. 1999).

### The KIX domain of Rex3 is required for 3’ Processing of RNase MRP RNA

Little is known about the cellular substrates of REXO1 and Rex3 proteins. Yeast Rex3 is required for the accurate 3’ end processing of the *NME1* transcript, the RNA component of RNase MRP, and functions redundantly with Rex1 and Rex2 in the processing of RNase P RNA, U5 snRNA and the autoregulated degradation of *RTR1* mRNA (Hodko et al. 2016; van Hoof et al. 2000). Studies in mammalian cells link the function of elongin A to transcription of enhancer RNAs and rRNA synthesis (Ardehali et al. 2021). To address the requirement for the N-terminal region of Rex3 for its function *in vivo*, we expressed full-length and N-terminal truncated versions of Rex3 as zz fusion proteins in yeast and assayed the expression levels of the fusion proteins and the profiles of *NME1* transcripts. Two *rex3* deletion constructs were generated, one which lacks the KIX domain (Δ1-106) and one which lacks both the KIX domain and the CHORD domain (Δ1-218) (Fig. 3E).

In agreement with earlier studies (van Hoof et al. 2000), the *rex3Δ* deletion strain accumulated MRP RNA transcripts that were longer than those seen in an isogenic wild-type strain. Expression of the wild-type Rex3 fusion protein restored the normal length of MRP RNA, showing that the fusion protein does not detectably affect the function of Rex3 (Fig. 3E). Deletion of the KIX domain increased the steady state expression level of Rex3 and blocked 3’ end processing of the RNase MRP RNA. Deletion of both the KIX domain and the CHORD domain led to a clear reduction in Rex3 protein levels and did not complement the processing phenotype of the *rex3Δ* mutant. We conclude that the N-terminal KIX domain within yeast Rex3 is required for its role in MRP RNA processing. Why loss of the KIX domain causes an increase in steady state levels of Rex3 is not clear but a plausible explanation is that Rex3 autoregulates its expression in a manner analogous to its role in the regulation of the *RTR1* mRNA (Hodko et al. 2016).

The CHORD domains of plant RAR1 proteins are protein interaction domains that mediate contacts with proteins containing a CS domain (Dubacq et al. 2002) such as the Hsp90 cochaperone Sgt1. One potential explanation for the depletion of the Rex3 mutant lacking the CHORD domain is that Rex3 requires interaction with heat shock proteins for its stable expression. Another potential Rex3 interactor is the yeast H/ACA snoRNP biogenesis factor Shq1, which contains a CS domain (Singh et al. 2009) that might couple Rex3-mediated snoRNA processing to snoRNP assembly. However, northern blot analyses did not reveal any changes in the lengths or relative abundance of the H/ACA snoRNAs snR8 or snR30 in the *rex3Δ* mutant, relative to the wild-type strain (Fig. 3E).

### Rex1 homologues are ubiquitous in eukaryotes while Rex3 proteins are restricted to fungi and animals

To address the evolutionary conservation of Rex1- and Rex3-related proteins, multiple sequence alignments and phylogenetic analyses were carried out on annotated domain sequences of eukaryotic DEDDh exoribonucleases. A phylogenetic analysis of the Rex1/Rex3 clade, together with a multiple sequence alignment of their Exo I, II and III motifs, is shown in Figure 4. The presence of the AlkP, RRM, KIX and CHORD domains within the Alphafold models of each protein is indicated on the right of the alignment.

**Figure 4.**
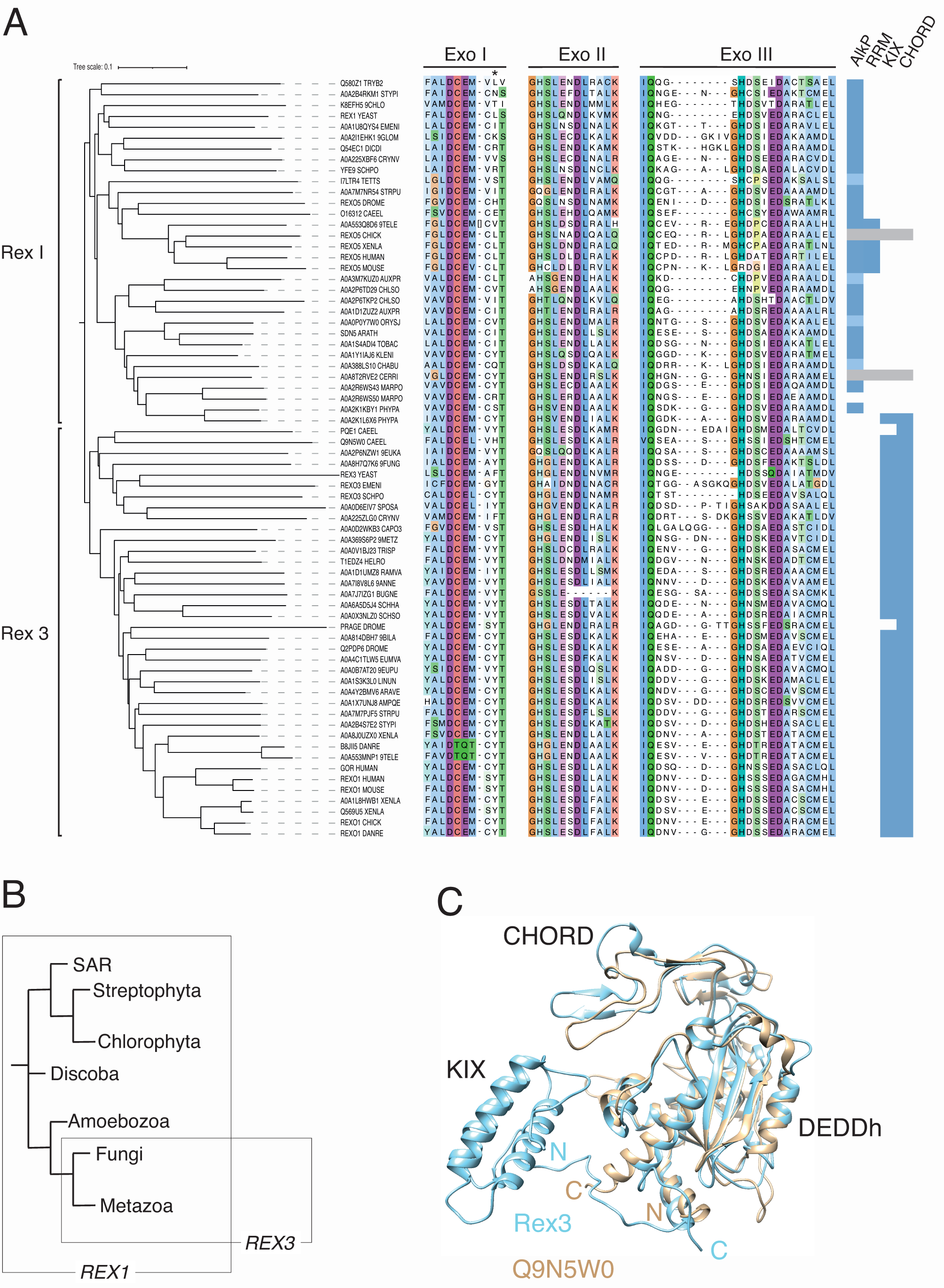
Phylogenetic analyses of Rex1 and Rex3 proteins. (A) Phylogenetic tree and multiple sequence alignment of Rex1 and Rex3 proteins. Exo I, II and III motif sequences are shown. The asterisk indicates a conserved Y/F residue in Rex3 proteins. AlkP, RRM, KIX and CHORD domains, determined by visual inspection of Alphafold structures, are indicated (light blue indicates partial or low confidence models; grey indicates the absence of the model). Proteins are identified by their common names or InterPro codes. Organism codes are as follows: TRYB2, *T. brucei*; STYPI, *S. pistillata*; 9CHLO, *Bathycoccus prasinos*; YEAST, *S. cerevisiae*; EMENI, *A. nidulans*; 9GLOM, *R. irregularis*; DICDI, *D. discoideum*; CRYNV, *C. neoformans*; SCHPO, *S. pombe*; TETTS, *T. thermophila*; STRPU, *S. purpuratus*; DROME, *D. melanogaster*; CAEEL, *C. elegans*; 9TELE, *D. translucida*; XENLA, *X. laevis*; AUXPR, *A. protothecoides*; CHLSO, *C. sorokiniana*; ORYSJ, *O. sativa*, subsp. *japonica*; ARATH, *A. thaliana*; TOBAC, *N. tabacum*; KLENI, *K. nitens*; CHABU, *C. braunii*; CERRI, *C. richardii*; MARPO, *M. polymorpha*; PHYPA, *P. patens*; EUKA, *Planoprotostelium fungivorum*; 9FUNG, *Umbelopsis vinacea*; SPOSA, *Sporidiobolus salmonicolor*; CAPO3, *Capsaspora owczarzaki*; 9METZ, *Trichoplax sp. H2*; TRISP, *Trichinella spiralis*; HELRO, *Helobdella robusta*; RAMVA, *Ramazzottius varieornatus*; 9ANNE, *Dimorphilus gyrociliatus*; BUGNE, *Bugula neritina*; SCHHA, *Schistosoma haematobium*; SCHSO, *Schistocephalus solidus*; 9BILA, *A. steineri*; EUMVA, *Eumeta variegata*; 9EUPU, *Arion vulgaris*; LINUN, *Lingula unguis*; ARAVE, *Araneus ventricosus*; AMPQE, *Amphimedon queenslandica*; DANRE, *D. rerio*, (B) Simplified cladogram showing the ancestry of *REX1* and *REX3* genes. (C) Structural overlay of Alphafold models for Rex3 from *S. cerevisiae* and Q9N5W0 from *C. elegans*. Domains and termini are labelled.

The Rex1/Rex3 family can be divided into two distinct clades on the basis of both sequence and structure criteria. Rex1-related proteins are found in all major eukaryotic lineages, including fungi, amoebae, metazoans, plants, green algae and in the protists *Trypanosoma brucei* (Discoba clade) and *Tetrahymena thermophila* (SAR clade). This suggests that the ancestoral *REX1* gene arose early during eukaryotic evolution. No Rex1 homologue was found for *Danio rerio* but a homologue was identifiable in the closely related cyprinid fish, *Danionella translucida.* Similarly, we did not identify a Rex1 homologue in the genomes of the green alga *Chlamydomonas reinhardtii* or the spikemoss *Selaginella moellendorffii* but homologous proteins were identified in the algae *Auxenchlorella protothecoides*, *Klebsormidium nitens* and *Chlorella sorokiniana*, as well as the stonewort *Chara braunii* and common liverwort, *Marchantia polymorpha*. *M. polymorpha* encodes two Rex1-related proteins, one that has a predicted structure typical of the Rex1 family and one that is essentially just the DEDDh domain. In four of the available thirty models, the AlkP domains were clearly recognisable but either partial or of low confidence (indicated with light blue shading in Fig. 4A). The *half-pint* twin RRM domain was restricted to Rex1 proteins from vertebrates. In contrast to an earlier report (Silva et al. 2017), the mouse Rexo5 protein has a histidine to arginine substitution within the Exo III motif (Fig. 4). This is not inconsistent with its inferred exonuclease activity, since DEDDh family members with noncanonical catalytic centres have been shown to have inherent exoribonuclease activity (Daugeron et al. 2001; Wagner et al. 2007).

The Rex3 clade includes proteins from fungi and the REXO1 proteins from metazoans but lacks proteins from plants or algae. The ancestral *REX3* gene therefore appears to have evolved before the divergence of fungi and metazoans, although a Rex3 homologue was found in the mycophagus amoeba *Planoprotostelium fungivorum*. The canonical domain architecture for Rex3-related proteins consists of a tandemly arranged CHORD domain and DEDDh domain at the C-terminus, with an upstream KIX domain. This domain architecture has previously been noted for a CHORD domain protein from the basidiomycete *Rhizopogon vesiculosus* (Kaur and Subramanian 2018). An exception is the protein from *Trichinella spiralis* and related species, which has a C-terminal thioredoxin domain. An unusual architecture is shown by the Rex3 homologue from the rotifer *Adineta steineri*, which has two tandem copies of the DEDDh domain at the C-terminus, each with an upstream CHORD domain. Rex1- and Rex3-related proteins typically contain the sequence DCEM within the Exo I motif, the dipeptide DL within the Exo II motif and the dipeptide IQ upstream of the Exo III motif (Zuo and Deutscher 2001). We noticed a very strong preference for a tyrosine or phenylalanine residue within the Exo I motif of Rex3 homologues (indicated with an asterisk in Fig. 4). One possibility is that this hydrophobic residue contributes to substrate binding and/or specificity. Some noncanonical catalytic centres are also observed in the Rex3-related proteins. B8JII5 from *D. rerio* and A0A553MNP1 from *D. translucida* have E/Q substitutions in the Exo I motif, while A0A7J7IZG1 from *Bugula neritina* lacks part of Exo II, including the conserved aspartate residue. Structural overlay of the Alphafold model for the *B. neritina* protein with that of yeast Rex1 suggests the overall fold of the catalytic centre is nevertheless conserved and that an electron pair could potentially be donated by either S840 or T843.

Notably, *D. melanogaster* and *Caenorhabditis elegans* express a Rex3 variant that lacks an N-terminal KIX domain (Prage and Q9N5W0_CAEEL, respectively). Prage has an N-terminal region of low structure propensity, whereas a high confidence Alphafold model of the Q9N5W0 protein suggests a compact domain structure comprising solely of the CHORD and DEDDh domains (Fig. 4C). Inspection of the structure similarity clusters within Alphafold suggest that this structural variant is widespread among metazoans and also found in nonpathogenic *Cryptococcus* fungi. Prage is required for turnover of maternally expressed mRNAs during early embryogenesis in flies (Cui et al. 2016; Tadros and Lipshitz 2005). Some eukaryotes, therefore, encode a functional variant of Rex3 that lacks the KIX domain, in addition to the canonical Rex3 protein.

This study reports structural differences that differentiate closely related Rex1- and Rex3 DEDDh ribonucleases and provides evidence that characteristic conserved domains and motifs within the *S. cerevisiae* enzymes are required for RNA processing functions *in vivo*. Given the high degree of sequence similarity between Rex1 and Rex3 proteins, compared with other DEDDh exoribonucleases, and the restriction of Rex3 to specific eukaryotic lineages, it is probable that the ancestral *REX3* gene evolved from *REX1*. These findings suggest numerous further lines of investigation into mechanistic studies on the structure/function relationship of these enzymes.

## Materials and Methods

### Bioinformatics Analyses

Search and sequence analyses were carried out using the EMBL-EBI tools services (Madeira et al. 2019). Initially, DEDDh family exoribonucleases were identified through homology with the DEDDh domain of the yeast family members by iterative PSI-BLAST searches against the UniProtKb swissprot database or retrieved directly from the Ensembl Databank resource (Martin et al. 2023). Retrieved sequences were limited to proteins from humans and the model organisms *A. thaliana, C. elegans*, *D. rerio, Dictyostelium discoideum, D. melanogaster, Gallus gallus, Mus musculus, S. cerevisiae*, *S. pombe* and *Xenopus laevis*. Additional proteins within InterPro entry IPR047021 (RNA exonuclease REXO1/REXO3/REXO4-like) were selected from the organisms *A. nidulans, A. protothecoides*, *Ceratopteris richardii, C. braunii, C. reinhardtii, C. sorokiniana*, *C. neoformans, D. translucida, K. nitens, M. polymorpha, Nicotiana tabacum, Oryza sativa* subsp*. japonica, Physcomitrium patens, Rhizophagus irregularis*, *S. moellendorffii, Stronglyocentrotus purpuratus, Stylophora pistillata, T. thermophila* and *T. brucei.* Further Rex3-related proteins were retrieved from the InterPro entry IPR031736 (RNA exonuclease 1 homolog-like domain) that also contained an annotated DEDD nuclease domain. Identical sequences were filtered using UniProt Align. Multiple sequence alignments were performed using MAFFT and viewed in JalView (Waterhouse et al. 2009). Phylogeny trees were generated using iTOL (Letunic and Bork 2021). Alphafold structural models (Jumper et al. 2021; Varadi et al. 2022) within listed structure similarity clusters (Barrio-Hernandez et al. 2023; Steinegger and Soding 2018) or accessed through InterPro entries were compared manually.

The following experimentally determined structures were downloaded from PDB (Berman et al. 2000): yeast Gal11/Med15 (PDB ID 2K0N) (Thakur et al. 2008); yeast TFIIS (PDB ID 3PO3) (Cheung and Cramer 2011); mouse CBP (PDB ID 4I9O)(Wang et al. 2013); human RECQL5 (PDB ID 4BK0)(Kassube et al. 2013); PUF60 with and without bound RNA (PDB IDs 5KW1, 5KWQ)(Hsiao et al. 2020); *E. coli* alkaline phosphatase (PDB ID 1ALK)(Kim and Wyckoff 1991) and iPGM from *B. stearothermophilus* (PDB ID 1EJJ) (Jedrzejas et al. 2000). Protein structure overlays and structure-based multiple sequence alignments were generated in UCSF Chimera (Pettersen et al. 2004). Alphafold structures were screened for matches within PDB and vice versa using the Dali server (Holm et al. 2023).

### Yeast Methods

*REX1 and REX3* coding sequences were amplified from yeast genomic DNA using the primers shown in Table 1 and cloned using standard cloning techniques into a plasmid that allows expression in yeast of N-terminal zz fusion proteins from the *RRP4* promoter (Mitchell et al. 1996). To generate the *rex1* deletion mutants, site-directed mutagenesis was performed on the wild-type expression construct using divergent primer pairs that introduce *Bam*HI sites and Gly/Ser linker sequences at appropriate positions. The *rex1* loop 1Δ mutant (ΔA145-S154) contains the linker GGSGSG, the loop 2Δ mutant (ΔN459-D472) contains the linker GSGSGSGSG and the loop 3Δ mutant (ΔR507-G546) contains the linker GSGSGG. The linker sequences were designed to be of minimal length yet sufficient to span the distance between residues at the beginning and ends of the loops, based on the Alphafold structural model. Constructs were validated by sequence analysis of the complete insert.

**Table 1.**
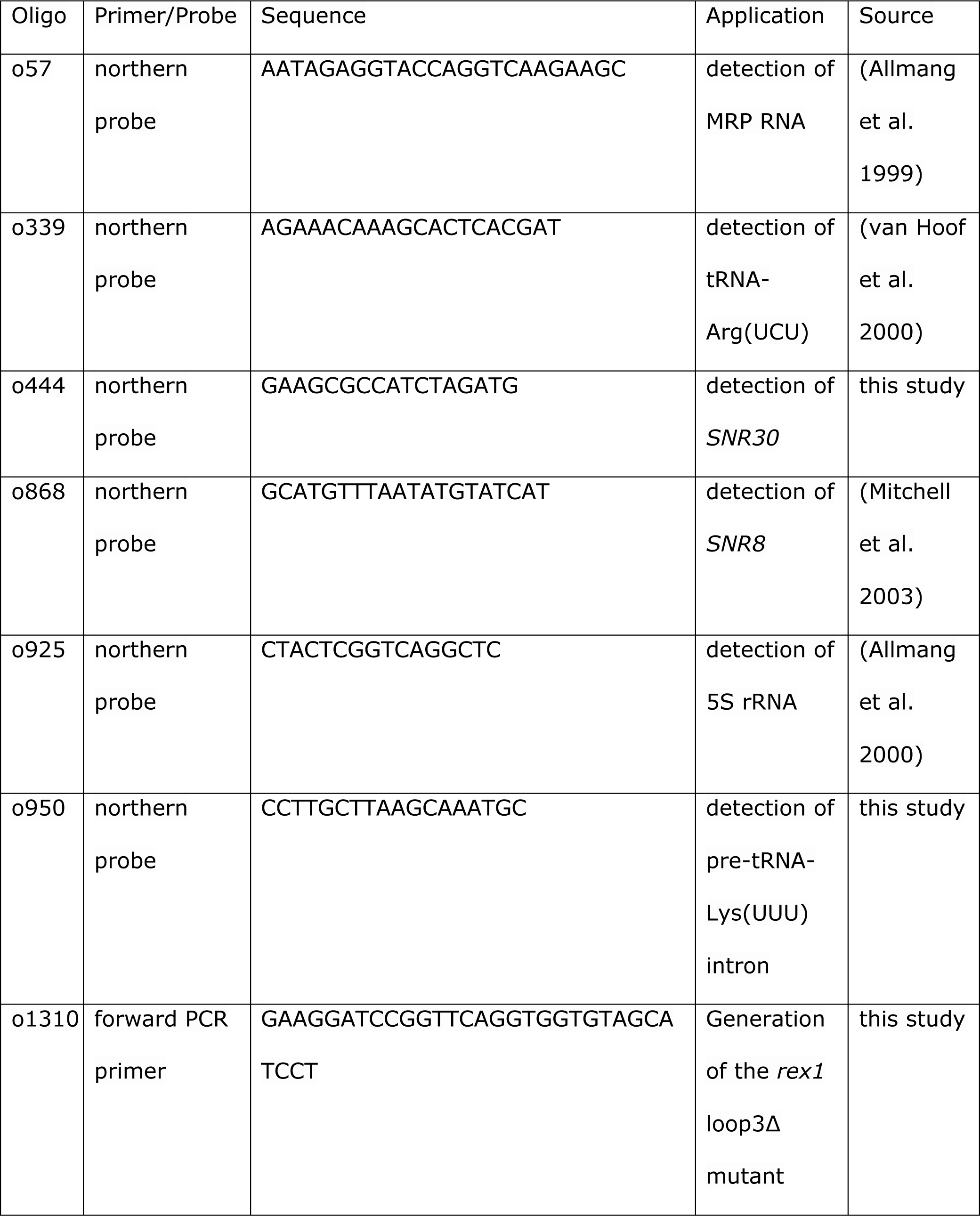
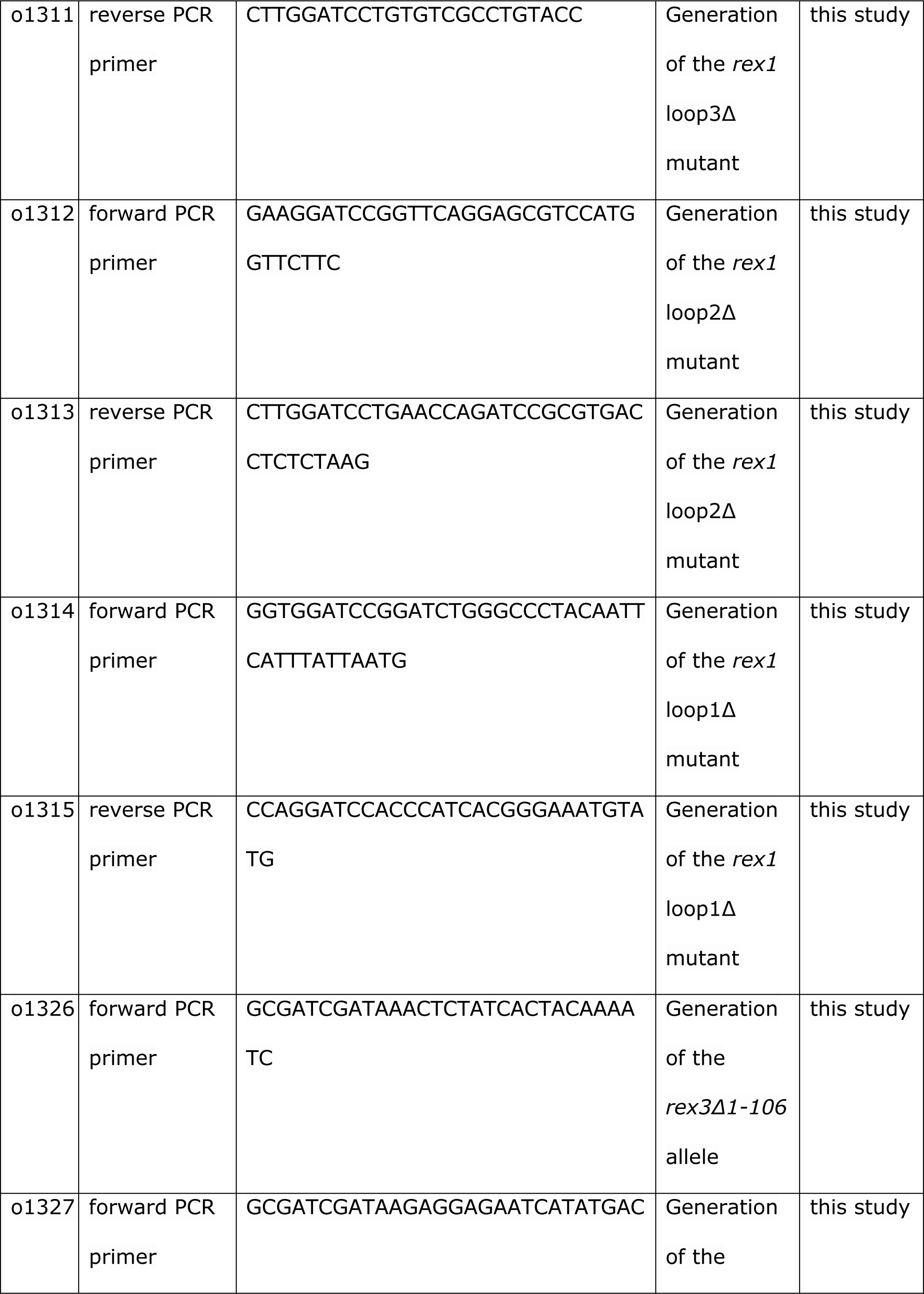
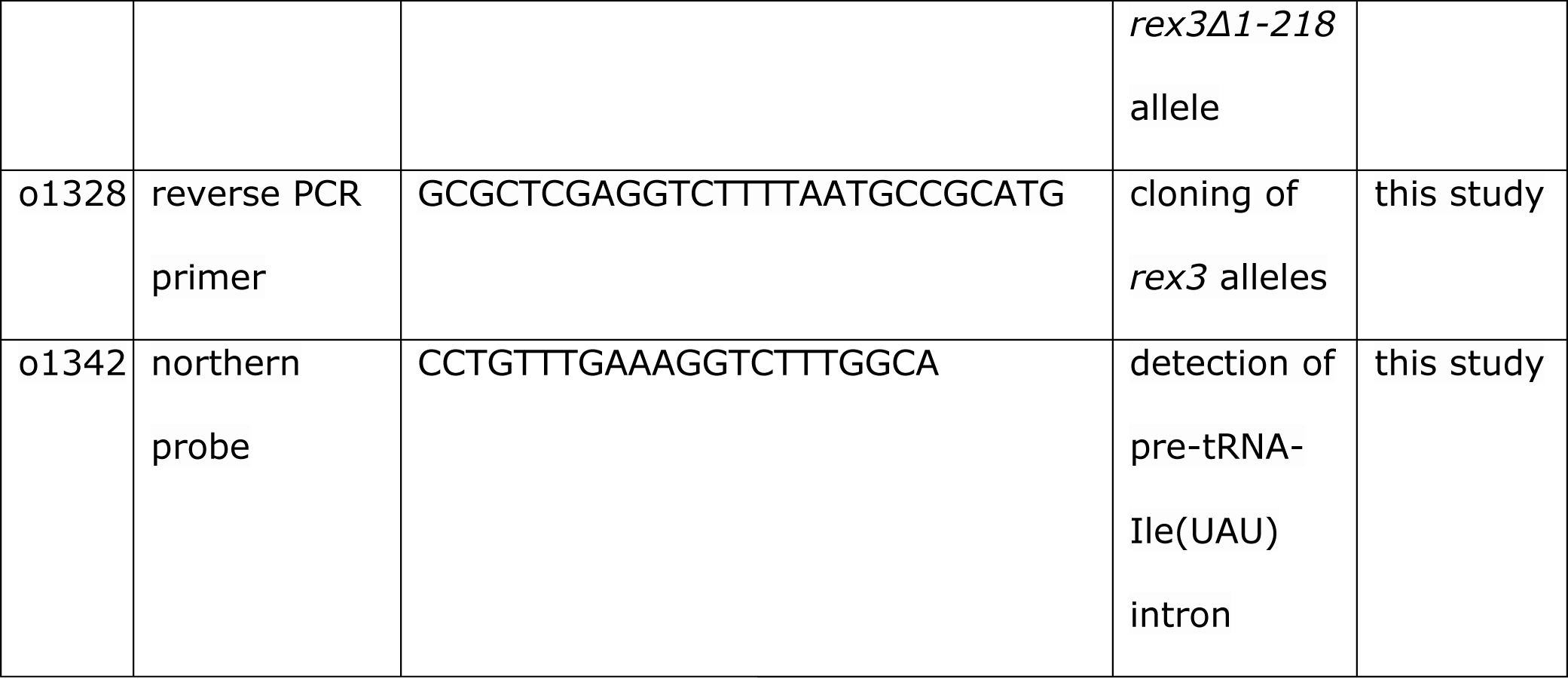
Oligonucleotides used in this study.

**Table 2.** Plasmids used in this study.

| Plasmid | Description | Source |
| --- | --- | --- |
| p674 | wild-type Rex1 expressed as an N-terminal zz protein from the *RRP4* promoter in pRS313. | (Daniels et al. 2022) |
| p966 | loop 2Δ derivative of p674 | this study |
| p967 | loop 1Δ derivative of p674 | this study |
| p972 | loop 3Δ derivative of p674 | this study |
| p973 | wild-type Rex3 expressed as an N-terminal zz protein from the *RRP4* promoter in pRS416. | this study |
| p974 | Δ1-106 variant of p973 | this study |
| p975 | Δ1-218 variant of p973 | this study |

Yeast strains were grown in selective SD minimal medium (2% glucose, 0.5% ammonium sulphate, 0.17% yeast nitrogen base, plus appropriate amino acids and bases) lacking uracil or histidine. Solid medium contained 2% bacto-agar. Plasmid transformations were performed using a standard lithium acetate procedure (Gietz et al. 1992). Yeast *rex1 rrp47Δ* double mutants were isolated by plasmid shuffling on medium containing 5’-fluoro-orotic acid. The plasmid shuffle strain (*Mat**a** ade2 ade3 his3 leu2 trp1 ura3 rex1Δ::KANMX4 rrp47Δ::KANMX4* + pRS416[*RRP47,ADE3*]) has been described previously (Costello et al. 2011). Spot growth assays were performed on ten-fold serial dilutions of saturated cultures from three independent FOA-resistant isolates of each *rex1 rrp47Δ* double mutant. Cells were grown on YPD medium at 30°C and photographed after 2 days. Yeast *rex1Δ::KANMX4* and *rex3Δ::KANMX4* strains in the BY4741 background (*Mata his3Δ1 met15Δ0 leu2Δ0 ura3Δ0*) were obtained from Euroscarf (Frankfurt, Germany).

### Protein and RNA Analyses

Denatured protein whole cell lysates were prepared under alkaline conditions, as described previously (Daniels et al. 2022). Western blot analyses were carried out using a peroxidase/antiperoxidase conjugate (P1291, Sigma Aldrich) to detect zz fusion proteins. Pgk1 was detected using a mouse monoclonal antibody (clone 22C5D8, Life Technologies) followed by an HRP-conjugated goat anti-mouse secondary antibody (1706516, Bio-Rad Laboratories). Proteins were visualised by ECL using an iChemi XL GelDoc system fitted with GeneSnap software (SynGene) and quantified using ImageJ (NIH). Rex1 and Rex3 expression levels were determined on three biological replicates, normalised to the expression level of Pgk1 and expressed relative to the level of the wild-type protein.

Total cellular RNA was prepared using a standard guanidinium hydrochloride/phenol extraction method. RNA was resolved through denaturing polyacrylamide/urea gels and transferred to Hybond N^+^ membranes (GE Healthcare, UK) using standard protocols. Northern hybridisations were carried out using 5’-^32^P-labelled oligonucleotides. ^32^P signals were captured using PhosphoImager screens and retrieved using a Typhoon FLA 7000 imager. RNA analyses were performed on two biological replicates.

## Acknowledgements

P.D. was supported by the White Rose BBSRC doctoral training programme and the BBSRC grant BB/V00722X/1. S.K. and I.T. were on MBiolSci undergraduate programmes within the University of Sheffield. The authors thank Monika Feigenbutz for critical reading of the manuscript.

